# Evidence that recurrent circuits are critical to the ventral stream’s execution of core object recognition behavior

**DOI:** 10.1101/354753

**Authors:** Kar Kohitij, Kubilius Jonas, Schmidt Kailyn, Elias B. Issa, James J. DiCarlo

## Abstract

Non-recurrent deep convolutional neural networks (DCNNs) are currently the best models of core object recognition; a behavior supported by the densely recurrent primate ventral stream, culminating in the inferior temporal (IT) cortex. Are these recurrent circuits critical to ventral stream’s execution of this behavior? We reasoned that, if recurrence is critical, then primates should outperform feedforward-only DCNNs for some images, and that these images should require additional processing time beyond the feedforward IT response. Here we first used behavioral methods to discover hundreds of these “challenge” images. Second, using large-scale IT electrophysiology in animals performing core recognition tasks, we observed that behaviorally-sufficient, linearly-decodable object identity solutions emerged ~30ms (on average) later in IT for challenge images compared to DCNN and primate performance-matched “control” images. We observed these same late solutions even during passive viewing. Third, consistent with a failure of feedforward computations, the behaviorally-critical late-phase IT population response patterns evoked by the challenge images were poorly predicted by DCNN activations. Interestingly, deeper CNNs better predicted these late IT responses, suggesting a functional equivalence between recurrence and additional nonlinear transformations. Our results argue that automatically-evoked recurrent circuits are critical even for rapid object identification. By precisely comparing current DCNNs, primate behavior and IT population dynamics, we provide guidance for future recurrent model development.

## INTRODUCTION

In a single, natural viewing fixation (~200 ms), primates can rapidly identify objects in the central visual field, despite various identity preserving image transformations, a behavior termed core object recognition (DiCarlo et al., 2012). Understanding the brain mechanisms that seamlessly solve this challenging computational problem has been a key goal of visual neuroscience (Riesenhuber and Poggio, 2000; Yamins and DiCarlo, 2016). Previous studies (Freiwald et al., 2009; Hung et al., 2005; Majaj et al., 2015) have shown that object categories and identities are explicitly represented in the pattern of neural activity in the primate inferior temporal (IT) cortex, and that specific IT neural population codes are sufficient to explain and predict primate core object recognition. Therefore, understanding how the brain solves core object recognition boils down to building a neurally-mechanistic (i.e. neural network) model of the primate ventral stream that, for any image, accurately predicts the neuronal firing rate responses at all levels of the ventral stream, including IT.

At present, the neural network models that best explain and predict the individual and population responses (image evoked, time averaged firing rates) of primate (macaque) IT neurons have been found in the architectural family of deep convolutional neural networks (DCNNs) trained on object categorization (Cadieu et al., 2014; Guclu and van Gerven, 2015; Yamins et al., 2014). These neural networks are also the current best predictors of primate behavioral patterns over dozens of core object recognition tasks (Rajalingham et al., 2018; Rajalingham et al., 2015). All neural networks in this model family are almost entirely feed-forward. Specifically, unlike the ventral stream (Felleman and Van Essen, 1991; Rockland et al., 1994; Rockland and Van Hoesen, 1994; Rockland and Virga, 1989), they lack cortico-cortical feedback circuits, sub-cortical feedback circuits, and medium to longrange intra-area recurrent circuits (as shown in Fig 1A). The short time duration (~200 ms) needed to accomplish accurate core object category and identity inferences in the ventral stream (Hung et al., 2005; Liu et al., 2009; Thorpe et al., 1996) suggests the possibility that recurrent-circuit driven computations are not critical for these inferences. In addition, it has been argued that recurrent circuits might operate at much slower time scales (Hinton et al., 1995), and thus may be much more relevant for processes like regulating synaptic plasticity to improve future behavior (learning). Taken together, the most parsimonious hypothesis is that core object recognition behavior does not require recurrent processing. The primary aim of this study was to try and falsify this hypothesis, and to provide new constraints to guide further neural network model development.

**Fig. 1.**
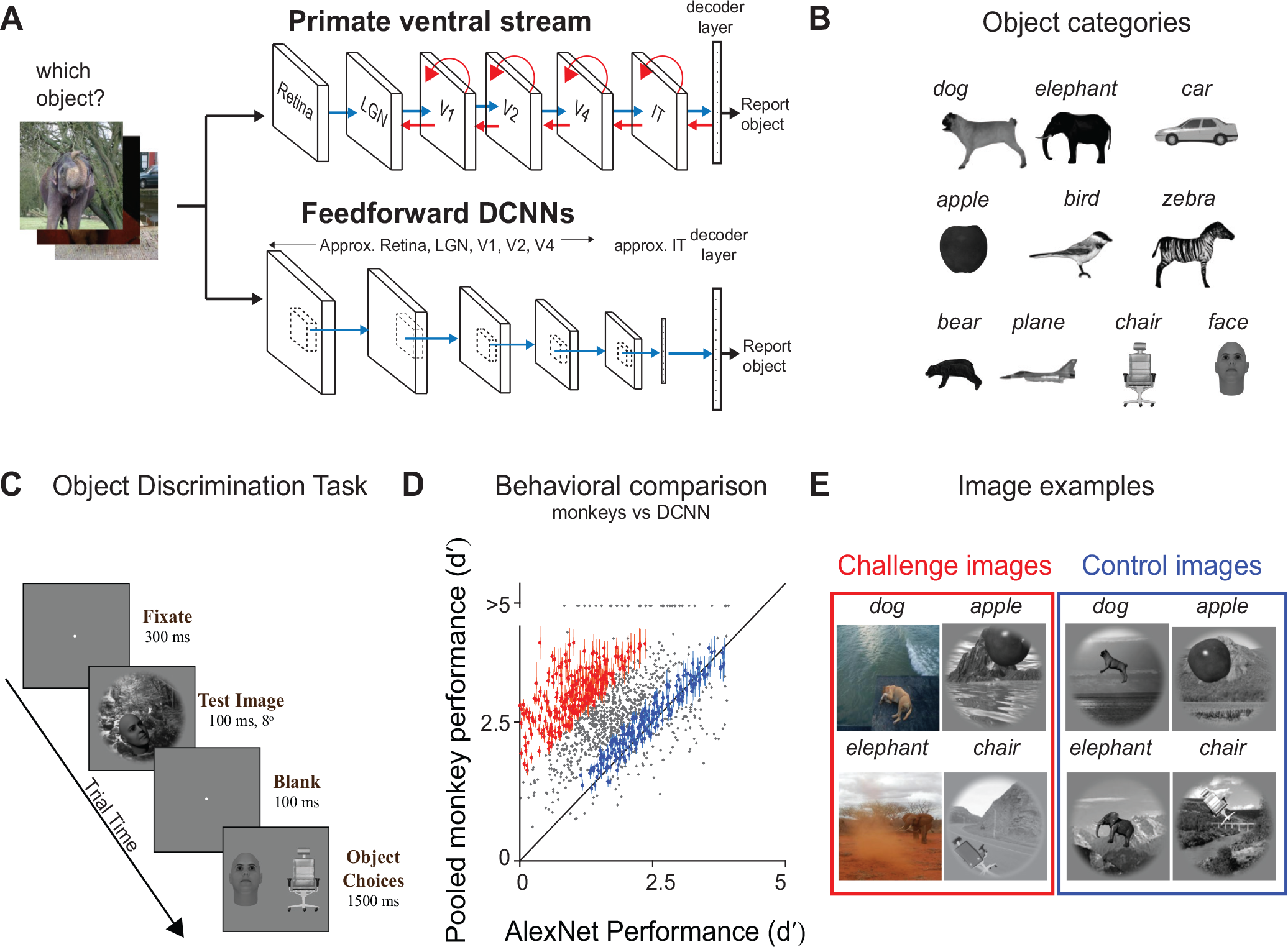
Behavioral screening and identification of control and challenge images. A) We task both primates (humans and macaques; top row) and feedforward DCNNs (bottom row) to identify which object is present in each Test image (1320 images). The top row shows the stages in the ventral visual pathway in primates (retina, LGN: lateral geniculate nucleus, areas V1, V2, V4, and IT), which is implicated in core object recognition. We can conceptualize each stage as rapidly transforming the representation of the image ultimately yielding to the primates’ behavior (i.e. producing a behavioral report of which object was present). The blue arrows indicate the known anatomical feedforward projections from one area to the other. The red arrows indicate the known lateral and top down recurrent connections. The bottom row demonstrate a schematic of a similar pathway commonly present in the DCNNs. These networks contain a series of convolutional and pooling layers with nonlinear transforms at each stage, followed by fully connected layers (which approximates macaque IT neural responses) that ultimately gives rise to the models’ simulated behavior. Note that the DCNNs only have feedforward (blue) connections. B) Object categories. We used ten different object types; bear, elephant, face, plane, dog, car, apple, chair, bird and zebra. C) Binary object discrimination task. Here we show the timeline of events on each trial. Subjects fixate a dot. The test image (8 deg) containing one of ten possible objects was shown for 100 ms. After a 100 ms delay, a canonical view of the target object (the one present in the test image) and a distractor object (from the other 9 objects). appeared, and the human or monkey indicated which object was present in the test image by clicking on or making a saccade to one of the two choices respectively. D) Comparison of monkey performance (pooled across 2 monkeys) and DCNN performance (AlexNet; ‘fc7’ Krizhevsky et al. 2012). Each dot represents the behavioral task performance (; refer Methods) for a single image. We reliably identified challenge (red dots) and control (blue dots) images. Error bars are bootstrapped s.e.m E) Examples of four challenge and four control images.

There is growing evidence that the feedforward DCNNs fall short of accurately predicting image-by-image primate behavior in a variety of situations (Geirhos et al., 2017; Rajalingham et al., 2018). We therefore hypothesized that specific images for which the object identities are difficult for non-recurrent DCNNs to solve, but are nevertheless easily solved by primates, might be critically benefiting from recurrent computations in the primates. Furthermore, previous research (for review see Lamme and Roelfsema, 2000) suggest that the impact of recurrent computations in the ventral stream should be most relevant at later time points in the image driven neural responses. Therefore we reasoned that IT neural population representations of objects in images in which those object inferences critically rely on the recurrent computations will require additional processing time to emerge (beyond the initial evoked IT population response that begins at ~90 ms; feedforward pass).

To discover such images, we behaviorally compared primates (humans and monkeys) and a particular non-recurrent DCNN (AlexNet ‘fc7’; Krizhevsky et al., 2012) to identify two groups of images — those for which object identity is easily inferred by the primate brain, but not solved by DCNNs (referred to here as “challenge images”), and those for which both primates and models easily infer object identity (referred to here as “control images”). To test our neural hypothesis, we simultaneously measured IT population activity in response to each of 1320 images, using chronically implanted multi-electrode arrays across IT cortex of both the left and right hemispheres of 2 monkeys, while monkeys performed an object discrimination task.

Our results revealed that object identity decodes from IT neural populations for the challenge images took an average of ~30ms longer to emerge (~145 ms from stimulus onset) compared to control images (~115 ms from stimulus onset). Consistent with previous results, we also found that the top layers of DCNNs optimized for object categorization performance predicted ~50% of IT image-driven neural response variance at the leading edge of the IT population response. However, this fit to the IT response was significantly worse (<20% explained variance) at later time points (150-200 ms post stimuli onset) — the time points where linear decoders show that the IT population solutions to these challenge images have emerged. Taken together, these results argue against feedforward only models for the brain’s execution of core object recognition, and instead imply a behaviorally-critical role of recurrent computations. Notably, we also found the same neural population phenomena while the monkeys passively viewed the images, implying that the putative recurrent mechanisms for successful core object inference in the primate are automatic and not strongly state or task dependent. Furthermore, we show that the observed image-by-image difference in DCNN and primate behavior along with precisely measured IT population dynamics for each image better constrain the next generation of ventral stream neural network models over previous qualitative approaches.

## RESULT

As outlined above, we reasoned that, if recurrent circuits are critical to core object recognition behavior, then current non-recurrent DCNNs should perform less accurately than the ventral stream for some images, and the first goal of this study was to discover many such “challenge” images. Rather than making assumptions about what types of images (occluded, cluttered, blurred, etc.) might most critically depend on feedback, we instead took a data driven approach to identify such images.

### Identification of DCNN challenge and control images

To compare the behavioral performance of primates (humans and macaques) and current DCNNs image-by-image, we used a binary object discrimination task that we have previously tested extensively (Fig 1C; Rajalingham et al., 2018; Rajalingham et al., 2015). For each trial, monkeys used an eye movement to select one of two object choices, after we briefly (100 ms) presented a test image containing one of those choice objects (see Primate Behavioral Testing in Methods). Once monkeys are trained in the basic task paradigm, they readily learn each new object over full viewing and background transformations in just one or two days and they easily generalize to completely new images of each learned object (Rajalingham et al., 2015). This suggests that this task taps into relatively natural visual behavior, and that the object learning is unlikely to produce strong changes in the ventral visual stream.

We tested a total of 1320 images (132 images of each of ten objects), in which the primary visible object belonged to one of 10 different object categories (Fig 1B). To make the task challenging, we included various image types (see Fig S1A): synthetic objects with high view variation (scale, position and rotation) on cluttered natural backgrounds (similar to the ones used in Majaj et al., 2015; Pinto et al., 2008), images with occlusion, deformation, missing object-parts, and colored photographs (MS COCO dataset; Lin et al., 2014).

Behavioral testing of all of these images was done in humans (n=88; Fig S2) and in monkeys (n=2; Fig 1D). We estimated the behavioral performance of the subject pool on each image, and that vector of image-wise *d′* values is referred to as *I*_1_ (see Methods; also refer Rajalingham et al., 2018). We collected sufficient data such that the reliability of the *I*_1_ vector was reasonably high (median split half reliability *ρ̃*, humans = 0.84 and monkeys = 0.88). To test the behavior of each DCNN model, we first extracted the image evoked features of the penultimate simulated neural layer, e.g. ‘fc7’ layer of AlexNet (Krizhevsky et al., 2012). We then trained ten linear decoders (see Methods) to derive the binary task performances, and used a different set of images to test each model. Fig 1D shows an image-by-image behavioral comparison between the pooled monkey population and AlexNet ‘fc7’. We identified “control” images (blue dots; Fig 1D) as those where the absolute difference in primate and DCNN performance do not exceed 0.4 (*d′* units), and we identified “challenge” images (red dots; Fig 1D) as those where the primate performance was at least 1.5 units greater than the DCNN performance. The object level data is elaborated in the panels of Fig S3.; Four examples each from both category of images are shown in Fig 1E. The challenge images were not idiosyncratic to our choice of the AlexNet (‘fc7’) model. Many of them also turned out to be challenge images for a range of other tested feedforward DCNNs, e.g., VGG-S (Chatfield et al., 2014; Sermanet et al., 2013), Zeiler and Fergus (2014); see Fig S1B.

Our results show that on average, both macaques and humans outperform AlexNet (‘fc7’). Most importantly, this image search procedure produced two groups of images: 1) 266 challenge images that are accurately solved by primates, but are not solved by a leading feedforward-only DCNN (AlexNet; but see later), and 2) 149 control images that are solved equally well by primates and the DCNN. On visual inspection, we did not observe any specific image property that differentiated between these two groups of images (analyzed in more detail below).

### Temporal evolution of image-by-image object representation in IT

Previous studies (Hung et al., 2005; Meyers et al., 2008) have shown that the identity of an object in an image is often accurately conveyed in the population activity patterns of the inferior temporal cortex in the macaque. Specifically, appropriately weighted linear combinations of the activities of these IT neurons can approximate how neurons in down-stream brain regions could integrate this information to form a decision about the object identity, and such weighted linear combinations can accurately predict the average behavioral performance in all tested core object recognition tasks (Majaj et al., 2015). In previous work, we assumed one weighted linear combination of the neural population response vector per object category (each is termed an “object decoder”), and we adopted that assumption here as well.

In this study, we aimed to compare and contrast these object decodes from IT for the challenge and control images. First, we wanted to know if these IT object decoders were as accurate as the primates for both types of images, as would be predicted from previous work (Majaj et al., 2015), as that would demonstrate that the ventral stream successfully solves the challenge images (images that are, by definition, not solved by current feedforward neural network models). Second, we reasoned that, if recurrent computations were crucial to these solutions, those computations would introduce additional processing time, and therefore IT object decodes for challenge images should emerge later than IT object decode for control images.

To estimate the temporal evolution of the IT object decode for each image, we used large scale multi-electrode array recordings (Fig 2A) to sample and record hundreds of neural sites across IT cortex in two awake, behaving macaques. In each monkey, we implanted three chronic 10 × 10 microelectrode arrays, inferior to the superior temporal sulcus (STS) and anterior to the posterior middle temporal sulcus (pMTS); each array sampled from ~25 mm^2^ of the posterior, central and anterior part of IT. Recording sites that yielded a significant visual drive 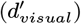, , high selectively and high image rank order response reliability 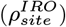 across trials were considered for further analyses (see Neural recording quality metrics in Methods). In total, we recorded from 424 valid IT sites which included 159 and 139 sites in the right hemisphere and 32 and 94 sites in the left hemisphere of monkey M (shown as inset in Fig 2A) and monkey N respectively.

**Fig. 2.**
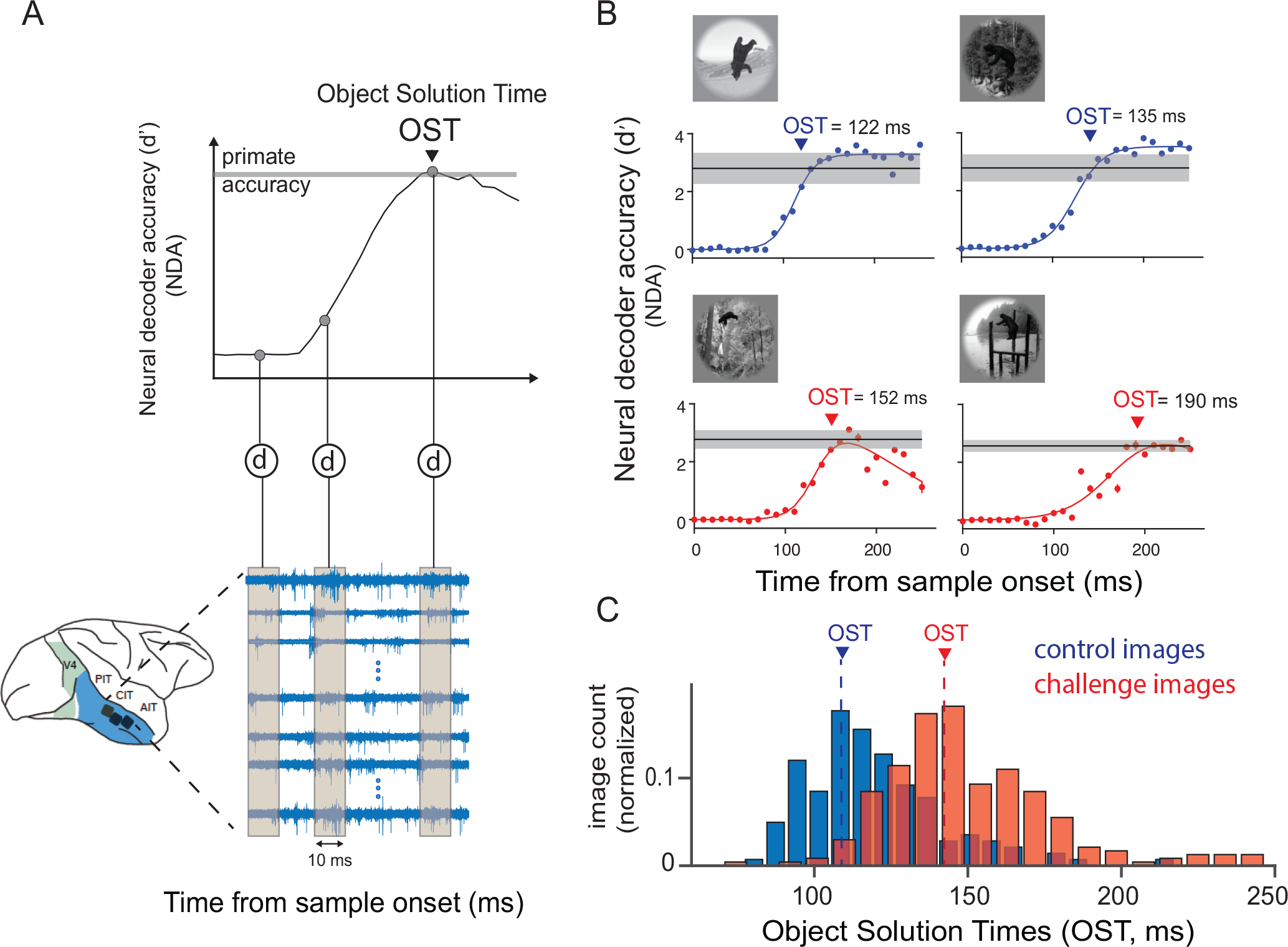
Large scale multi-unit array recordings in the macaque inferior temporal cortex. A) Schematic of array placement, neural data recording and object solution time estimation. We recorded extracellular voltage in IT from two monkeys, each hemisphere implanted with 2 or 3 Utah arrays. For each image presentation (100 ms), we counted multiunit spike events (see Methods for details), per site, in non overlapping 10 ms windows, post stimulus onset to construct a single population activity vector per time bin. These population vectors (image evoked neural features) were then used to train and test cross-validated linear support vector machine decoders (d) separately per time bin. The decoder outputs per image (over time) were then used to perform a binary match to sample task, and obtain neural decode accuracies (NDA) at each time bin. An example of the neural decode accuracy over time is shown in the top panel. The time at which the neural decodes equal the primate (monkey) performance, is then recorded as the object solution time (OST) for that specific image. B) Examples of IT population decodes over time, with the estimated object solution times for four images; two control (top panel: blue curves) and two challenge images (bottom panel: red curves). The red and blue dots are the estimated neural decode accuracies at each time bins. The solid lines are nonlinear fits of the decoder accuracies over time (see Methods). The gray lines indicate the performance of the primates (pooled monkey) for the specific images. Errorbar indicates bootstrapped s.e.m. C) Distribution of object solution times for both control (blue) and challenge (red) images. The median OST for control (blue) and challenge (red) images are shown in the plot with dashed lines.

To determine the time at which explicit object identity representations are sufficiently formed in the IT population activity, we plotted the trajectory of the IT object decode accuracy for each image and determined the time that it reached the level of the subject’s (pooled monkey) behavioral accuracy on that image. We termed this time, the “object solution time” (OST), and we emphasize that each image has a potentially unique solution time (*OST*_*image*_). Briefly, object solution time for each image, was defined as the time (relative to image onset) when the linear IT population decode (see Methods; Fig 2A, top panel) first rises to within the error margins of the pooled monkey behavioral score for that image. Because we recorded many repetitions of each image, we were able to measure the *OST_image_* very accurately (average standard error of ~9ms across all images, bootstrapped across repetitions).

Fig 2C shows the temporal evolution of the IT object decode and the OST estimates for two control images and two challenge images. For all four images, the correct answer is the object ‘bear’ (insets in Fig 2B). Two observations are apparent in these example images. First, for both the control and the challenge images, the IT decodes achieve the behavioral accuracy of the monkey (note, behavioral accuracy is similar for all four images, by design). Second, the IT decode solutions for challenge images emerge later than the solutions for the control images.

Both of these observations were also found on average in the full sets of challenge and control images. First, the IT decodes achieved the primate behavioral level of accuracy on average for the challenge and control imagesets (~91 % of challenge images and ~97 % of control images), which meant that we could determine an OST for all of these images. Second, and consistent with our hypothesis, we observed that IT object solution times (*OST*_*image*_) for the challenge images were, on average, ~30 ms later compared to the control images. Specifically, the median OST for the challenge images was 145±1.4 ms (median ± SE) from stimulus onset and the median OST for the control images was 115±1.4 ms (median±SE) (Fig 2C). Both of these results are consistent with the hypothesis that recurrent circuit computations are critical to core object recognition (see Introduction). Thus we next carried out a series of controls to rule out alternative explanations for these results.

### Comparison of initial visual drive in IT evoked by control and challenge images

We considered the possibility that the observed time lag for the challenge images’ OSTs might have been due to the IT neurons taking longer to start responding to these images. For example, if the information in those images took longer to be transmitted by the retina. However, the data do not support this possibility. First, we observed that control and challenge images share the same population neural onset response latencies (latency = 0.17 0.21 ms, median SE; paired t-test; t(423) = 0.3896, p = 0.69; see Fig 3A and Fig S4B), suggesting that the initial visual drive for the images in both sets arrive at approximately the same time in IT. However, we found that firing rates (R) were significantly higher (%R = 17.3%, paired t-test; t(423) = 6.8848, p <0.0001) for challenge images compared to control images, tested on a 30 ms window centered at 150 ms post stimuli onset. We do not yet know how to interpret this higher firing rate, but one possible explanation of this difference in IT mean firing rate is the effect of additional inputs from activated recurrent circuits into the IT neural sites at later time points (see Discussion). Regardless, these observations show that the challenge images drive IT neurons just as well as the control images during the early phase of the response.

**Fig. 3.**
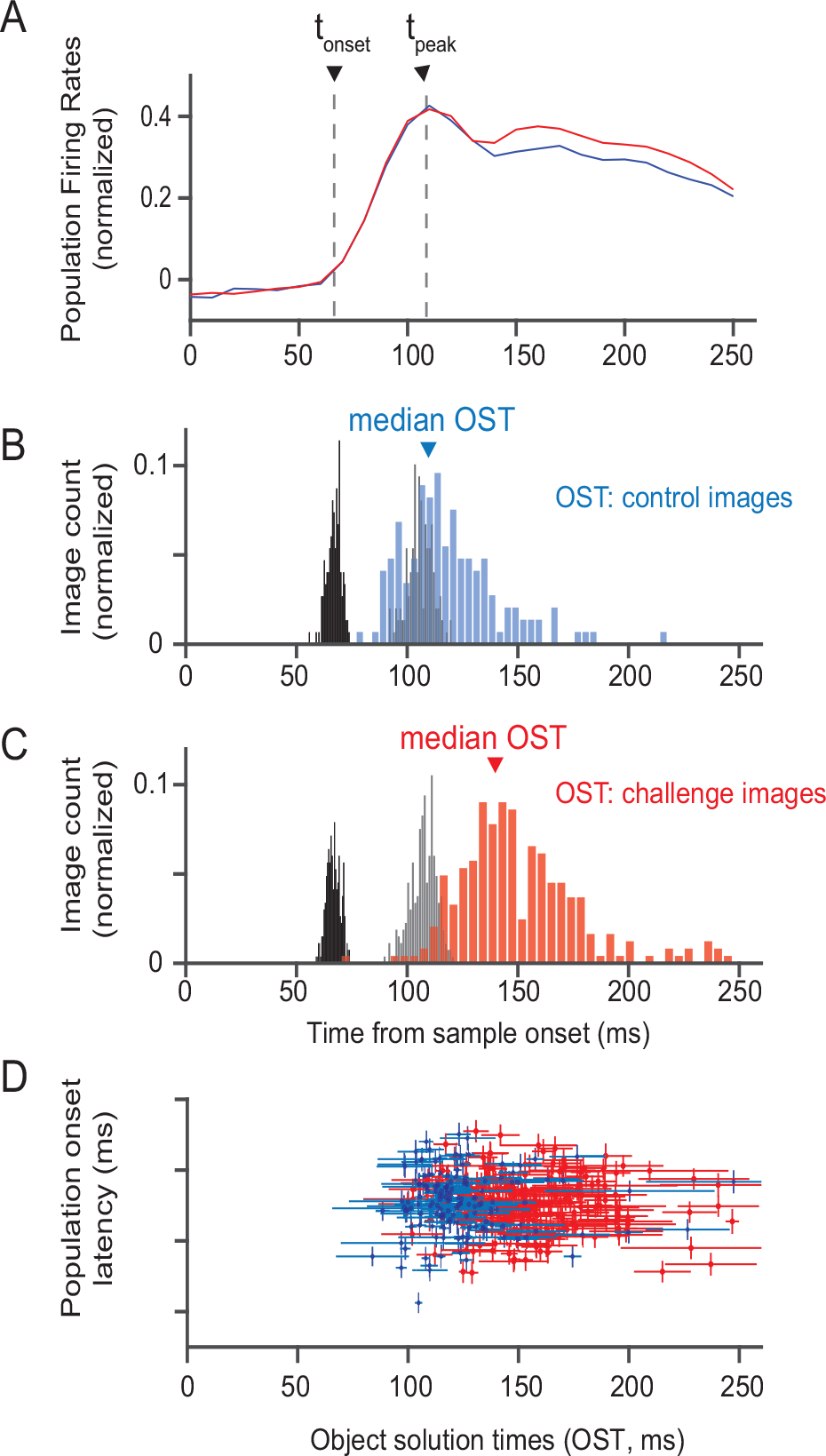
Relationship between object solution times and neural response latencies. A) Comparison of neural responses evoked by control (blue) and challenge (red) images. We estimated two measures of population response latency: Population onset latency (tonset) and Population peak latency (tpeak). B) Distributions of the population onset latencies (median across 424 sites), population peak response latencies (median across 424 sites) and object solution times for control images (n=149). C) Same as in B) but for challenge images (n = 266)D) Comparison of population onset latencies and object solution times for both control (blue) and challenge images (red). Vertical error bars show s.e.m across neurons and horizontal error bars show bootstrap (across trial repetition) standard deviation of object solution time estimates.

When we closely examined the neural population response latencies for each image, we found that the time at which the IT population firing rates started to increase from baseline (onset latency; t_*onset*_) and when the population firing rate reached its peak (t_*peak*_) did not coincide with the object solution times for that image (Fig 3B and 3C). We also found no correlation (Pearson r = 0.009; p = 0.8) between the population response onset latency for each image (see Methods) and the OST for that image (see Fig 3D). For example, inspection of Fig 3D reveals that some of the challenge images evoke faster-than-average latency responses in IT, yet have late OSTs (~200 ms). Conversely, some of the control images evoke slower-than-average IT responses, yet have very relatively fast OSTs (~110 ms). In sum, these results show that visual drive rapidly reaches IT for nearly all of these images, but that, for some images (mostly the challenge images), that visual driven population activity takes longer to evolve to an accurate, linearly-decodable format (OST).

### Comparison of effects of low level image properties on challenge and control image OSTs

We considered the possibility that the time lag for the challenge image OSTs might have been due to low-level image property differences between the two image-sets. From previous research, we know that temporal properties of IT neurons depend critically on low level image features like total image contrast energy (Oram, 2010), spatial frequency power distribution (Rolls et al., 1985), and spatial location of the visual objects (Op De Beeck and Vogels, 2000). So we asked if these low level explanations might explain the lag of the challenge image OSTs. First, we did not notice any significant differences (t_*onset*_ = 0.17 ms, paired t-test; t(423) = 0.3896; p=0.697) in neural firing rate onset latencies (Fig 3,, Fig S2B) between control and challenge images across the recorded neural sites. We also observed that solution times were not significantly correlated with image contrast (r = −0.04; p = .47). Second, we used the SHINE (spectrum, histogram, and intensity normalization and equalization) technique (Willenbockel et al., 2010) to equate low level image properties across the control and challenge image-sets, and re-ran the recording experiment (subsampling 118 images each from the control and challenge imagesets; no. of repetitions per image = 44; see Methods). The average estimated difference in OST values between “SHINED” challenge and control images was still ~24 ms (Fig S4C).Third, we sub-sampled the two image sets to make sure that the distribution of object eccentricity was perfectly matched between the two sets, but we still found an almost identical average OST lag between these two images sets (average ∆OST = 27 ms).

### Object solution estimates and timing during passive viewing

To test whether the late-emerging object solutions in IT are task dependent, we also recorded IT population activity during passive viewing of both the challenge and the control images. Monkeys fixated a dot, while images were each presented for 100 ms (same duration as the active task viewing of the image, see Figure. 1), followed by 100 ms of no image, followed by the next image for 100 ms, etc. (typically 5 images were presented per fixation trial; see Methods). A-priori, several outcomes of switching from active to passive viewing seemed likely: a decreased goodness of both the early-emerging and the late-emerging IT decoded solutions, a decreased goodness of the late-emerging solutions, a further delay in the late-emerging solutions, or no effect.

First, similar to the active condition (%∆R = 17.3%), we observed that challenge images evoked a significant higher firing rate (%∆R = 13.2%, paired t-test; t(423) = 8.27, p <0.0001) at later time points (tested on a 30 ms window centered at 150 ms post stimuli onset) compared to the control images. Second, similar to the active viewing, we observed that we could successfully estimate the object solution times for 92% of challenge and 98% of control images. The object solution times estimated during the active and passive conditions were also significantly correlated (Spearman r = 0.76; p <0.0001). Similar to the active condition, challenge image solutions, on average, required an additional time of ~28 ms to achieve full solution compared to the control images. In sum, we observe that the solutions in IT emerge with a similar lag and overall accuracy (goodness) during passive viewing. Therefore, we conclude that the putative recurrent signal that emerges in IT is not entirely task dependent and the mechanisms giving rise to this activity is fairly reflexive. This is consistent with previous findings of McKee et al. (2014), where they reported that IT cortex predominantly shows task-independent visual feature representation.).

### IT predictivity across time using current feedforward deep neural network models of the ventral stream

We reasoned that, if the late-emerging IT population solutions are indeed dependent on recurrent computations that are lacking in current DCNN models, then perhaps the previously demonstrated ability of those models to (partially) explain and predict individual IT neurons (Yamins et al., 2014) was due mostly to their ability to capture the feedforward portion of the IT response. To test this idea, we asked how well AlexNet features could predict the time-evolving IT neural population response vector up to and including the object solution time for each image. To do this, we used previously described methods. Specifically, we quantified that IT population goodness of fit as the median (over neurons) of the noise corrected explained response variance score (IT predictivity; Figure S4A; also see Methods; similar to Yamins et al., 2014).

First, we observed that the top layers (penultimate) of a performance optimized feedforward DCNN (AlexNet ‘fc7’) predict 44.3 ± 0.7 % of the potentially explainable IT neural response variance during the early phases (90-110 ms) of IT responses (Fig 4A) for all images This result provides further confirmation that feedforward DCNNs indeed approximate the early (putative largely feedforward) IT population response pattern. However, we observed that the ability of AlexNet ‘fc7’ model features to predict the IT population vector significantly worsened (<20% explained variance) as that response vector evolved over time(>150ms) for images with late OSTs (Fig 4A). This drop in IT predictivity was not due to low signal to noise ratio of the neural responses during those time points because our explained variance measure already compensates for any changes in SNR, and also because SNR remains relatively high in the late part of the IT responses (Fig S6). In sum, the gradual drop in IT predictivity by these feedforward DCNN models is consistent with the hypothesis that late-phase IT population responses are modified by the action of recurrent circuits that are not contained in those models. Consistent with our hypothesis that challenge images rely more strongly on recurrence than control images, we observed that the drop in IT predictivity coincided with the solution times of the challenge images (refer top panel histograms for OST distributions of challenge and control images).

**Fig. 4.**
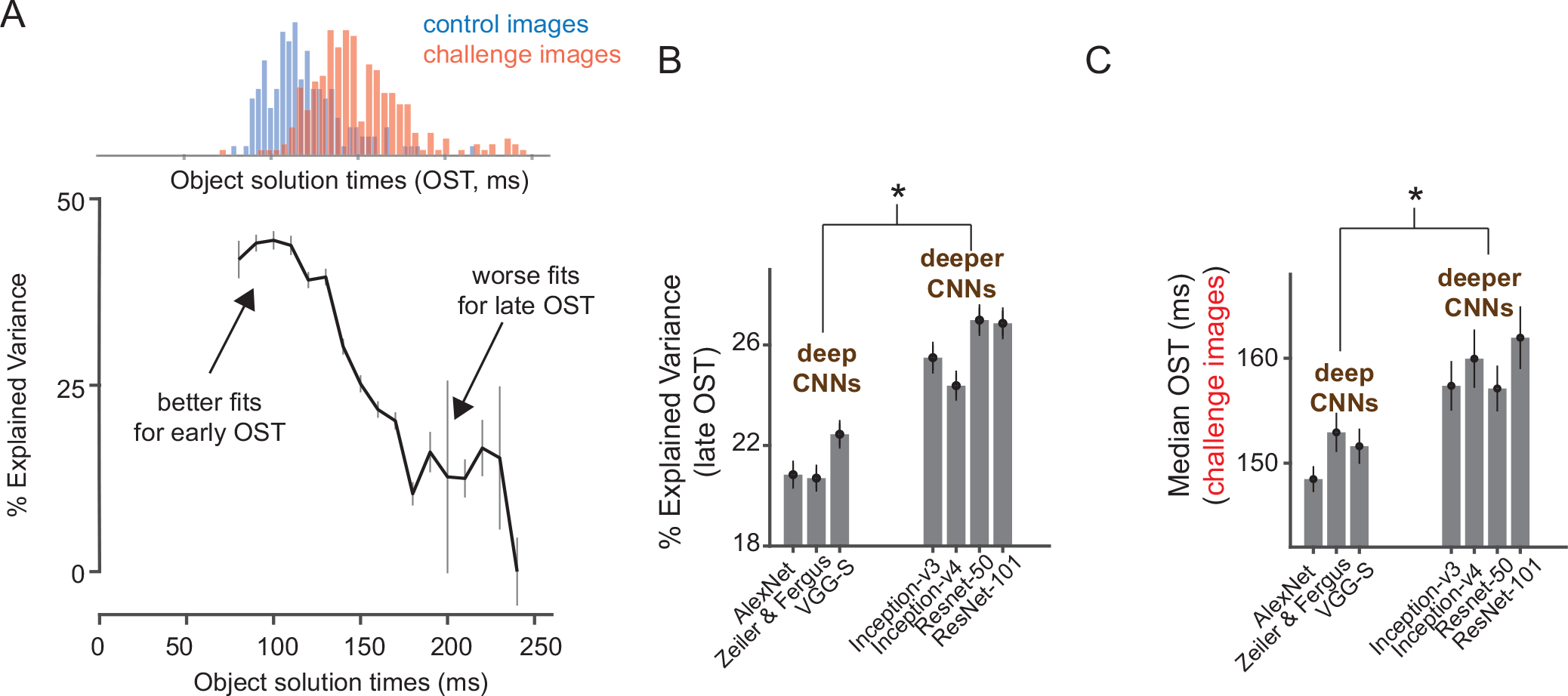
Predicting IT neural responses with DCNN features. A) IT predictivity of AlexNet’s ‘fc7’ layer as a function of object solution time (ms). For each time bin, we consider IT predictivity for images that have a solution time equal to or higher than that time bin. Error bars indicate the standard error of mean across neurons. Top panel shows the distribution of object solution times for control (blue) and challenge (red) images. B) IT predictivity computed separately for late OST images (OST>150 ms; total of 349 images) at the corresponding object solution times, as function of deep (AlexNet, Zeiler and Fergus, VGG-S) and deeper (Inception, ResNet) CNNs. * indicates a statistically significant difference between the two groups. C) Comparison of median OST for challenge images across CNNs of different depth. Images that remain unsolved (i.e. categorized by us as “challenge” images) by deeper CNNs showed even longer OSTs in IT cortex than the original full set of challenge images. * indicates a statistically significant difference between the two groups.

### Evaluation of deeper CNNs as models of ventral visual stream processing

Although, the above results suggest the likely importance of recurrent computations in the primate ventral stream for some images, we are still left with the open question: what specific computational role does recurrent circuits provide beyond the feedforward representation during core object recognition behavior? We reasoned that recurrence during core object recognition in the ventral stream is functionally equivalent to stacking further non-linear transformations onto the initial evoked (~feedforward) IT population response pattern.Therefore, we speculated that simulated neurons from deeper CNNs (those with a higher number of stacked nonlinear transformations) might better approximate the recurrent computations of the ventral stream, even though they were not specifically designed to emulate the many anatomical recurrent circuits of the ventral stream. To test this idea, we asked if existing very deep CNNs provide a better neural match to the IT response at its late phase and to the image-by-image patterns of behavioral performance. Currently there are many deeper CNNs available that outperform the baseline DCNN used here (AlexNet), such as inception-v3 (Szegedy et al., 2016), inception-v4 (Szegedy et al., 2017) and ResNet-50, ResNet-101(He et al., 2016). Based on the number of layers, we divided the tested DCNN models into two groups, deep (8 layers; AlexNet, Zeiler and Fergus model, VGG-S) and deeper (>20 layers, inception-v3, inception-v4, ResNet-50, ResNet-101) CNNs. We made three observations, that corroborate our speculation.

First, we searched all the above mentioned neural networks to determine which layers of the models best predict the late IT responses for the images with late OSTs. Interestingly, we observed that layers of deeper CNNs predict IT neural responses at the late phases significantly higher (%∆Predictivity = 10.72%, , paired t-test; t(423) = 8.36, p <0.0001) than shallower models like AlexNet (Fig 4B). This observation suggests that deeper CNNs might indeed be approximating “unrolled” versions of the ventral stream’s recurrent circuits. Second, as expected from the Imagenet challenge results (Russakovsky et al., 2015), we observed an increased performance and therefore reduced number of “challenge” images for deeper CNNs. Third, we found that the images that remain unsolved (i.e. categorized by us as “challenge” images) by these deeper CNNs showed even longer OSTs in IT cortex than the original full set of challenge images (Fig 4C). Assuming that longer OST is a signature of more recurrent computations, this suggests that newer, deeper CNNs have implicitly, but only partially, approximated — in a feedforward network — some of the computations that the ventral stream implements recurrently to solve some of the challenge images.

### Comparison of backward visual masking between challenge and control images

So far we have observed that feedforward DCNNs poorly predict the IT neural responses at later times beyond the putative feedforward response (90-110 ms post image onset), during which a majority of the challenge images (~82 %) develop their object solutions in IT. Based on these results, we hypothesized that these later IT population responses are critical for successful core object recognition behavior for the challenge images. To further test this idea, we performed an additional behavioral experiment that aimed to corroborate the neurophysiology results. We modified the original object discrimination task by adding a visual mask (phase scrambled image; Stojanoski and Cusack, 2014) for 500 ms (Fig 5A), immediately following the test image presentation: a paradigm commonly known as backward visual masking. This type of backward masking has been previously associated with selective disruption of the recurrent inputs to an area from other areas (Fahrenfort et al., 2007; Lamme et al., 2002). Given that solutions for the challenge images can arise in IT cortex only at later time points compared to the control images, we reasoned that if disruption in processing produced by a visual mask affects IT at earlier times, it will produce larger behavioral deficits for challenge images compared to control images. However, we predicted that these differences should subside at longer presentation times when enough time is provided for the recurrent processes to build a sufficient object representation for both control and challenge images in IT. Therefore, during this experiment, we tested a range of masking disruption times by randomly interleaving the sample image duration (and thus the mask onset). Specifically, we tested 34, 67, 100, 167 and 267 ms (see Methods). Our results (Fig 5B) show that visual masking indeed had a significantly stronger effect on the challenge images at smaller presentation durations compared to the control images. Consistent with our hypothesis, we did not observe any measurable masking differences between the two imagesets at longer presentation times (~267 ms). Median *d′* (difference between control and challenge images grouped by objects) averaged across all 10 objects were 0.5, 0.81, 0.33, 0.40, and −0.02 for 34, 67, 100, 167 and 267 ms presentation duration respectively. The difference in performance was statistically significant at the .05 significance level (Bonferronni adjusted) for all presentation durations except 267 ms. Together with the neurophysiology results, these observations provide converging evidence towards a critical role of recurrent ventral stream computations in the brain’s ability to infer object identity in the challenge images.

**Fig. 5.**
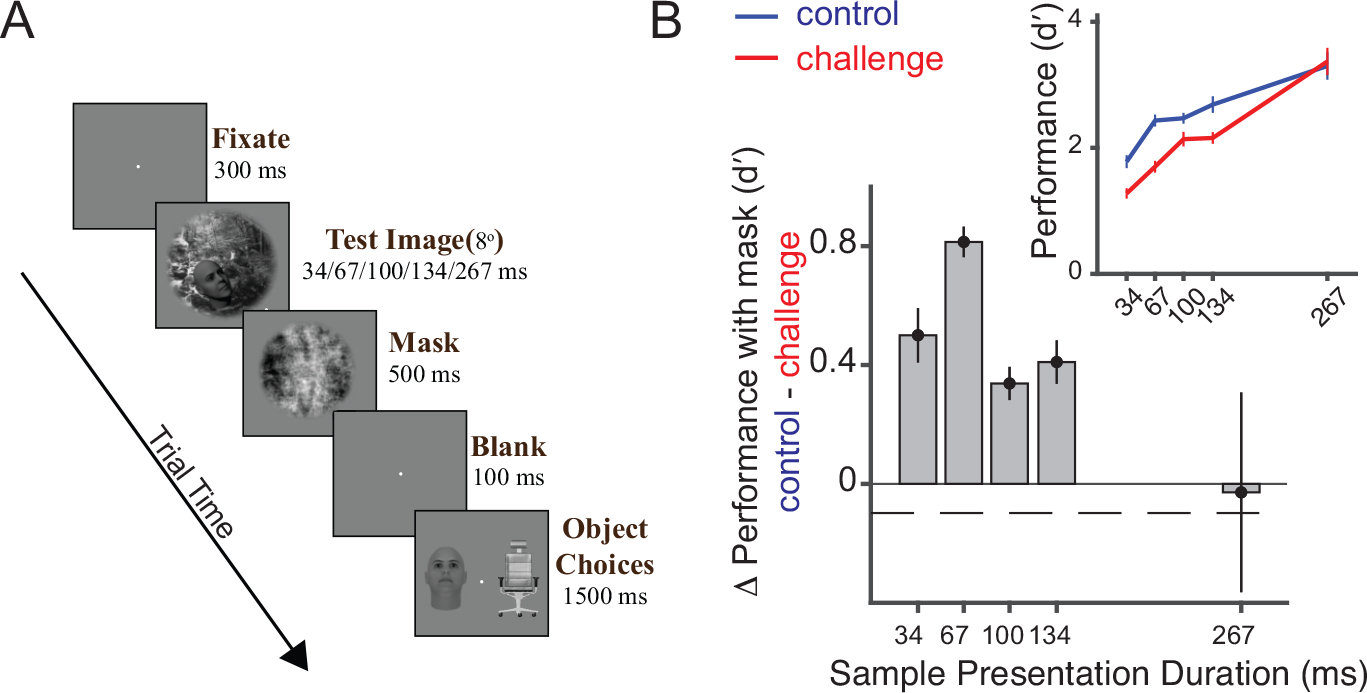
A) Binary object discrimination with backward visual masking. The test image (presented for 34, 67, 100, 134 or 267 ms) was followed immediately by a visual mask (phase scrambled image) for 500 ms, followed by a blank gray screen for 100 ms and then the object choice screen. Monkeys reported the target object by fixating it on the choice screen. B) Difference in behavioral performance between control and challenge image after backward visual masking. Each bar on the plot (y-axis) is the difference in the pooled monkey performance during the visual masking task (A) between control and challenge images at the respective sample image presentation durations (x-axis). The top panel inset shows the raw performance (*d′*) for the two groups of images (blue: control images, red: challenge images). Error bars denote the s.e.m. across all objects.

### Constraints for future models provided by our data

The results above motivate a change in the architecture of artificial neural networks that aim to model the ventral visual stream (i.e. addition of recurrent circuits) — motivating a switch from largely feedforward DCNNs to recurrent DCNNs. However, a primary goal of experiments is not simply to provide motivation, but to also provide validation and strong constraints for guiding the construction of new models. The results obtained here provide three precisely measured constraints for next generation neural network models. First, we provide a behavioral vector, ∆*d′* that quantifies the performance gap between current feedforward DCNNs (e.g. AlexNet) and the image-by-image primate core object recognition behavior (*I*_1_). For each of these images, we have estimated the time at which object solutions are sufficiently represented in the macaque IT cortex. So the other two constraints include the neural responses to each of the tested images at the object solution times as well as a vector of estimated OSTs (each element of the vector corresponds to an image). Next generation dynamic models of the ventral stream should be constrained to produce the target features (object solutions) at these times.

### Model-driven versus image-property driven approaches to study recurrence

Previous research has suggested that recurrent computations in the ventral stream might be necessary to achieve pattern completion when exposed to occluded images (Spoerer et al., 2017; Tang et al., 2017; Walther et al., 2005), object based attention in cluttered scenes (Bichot et al., 2015; Walther et al., 2005) etc. Indeed, we observe that several image properties like object size, presence of occlusion, and object eccentricity (Fig 6) are significant predictors (see Methods: Estimation of the OST prediction strength) of our putative recurrence signal (the *OST* vector). However, our results above suggest another possible image-wise predictor of ventral stream recurrence — the difference in performance between feed-forward DCNNs and primates, ∆*d′* (Fig 6). This vector is likely itself dependent on a complex combination of image properties, such as those mentioned above. However, it is directly computable and our results show that it can serve as a much better model guide. In particular, we find that ∆*d′* is significantly correlated with IT *OST* (Spearman *ρ* = 0.44; p < 0.001), and, in this sense, is a much better predictor of likely recurrence than any of the individual image properties.

**Fig. 6.**
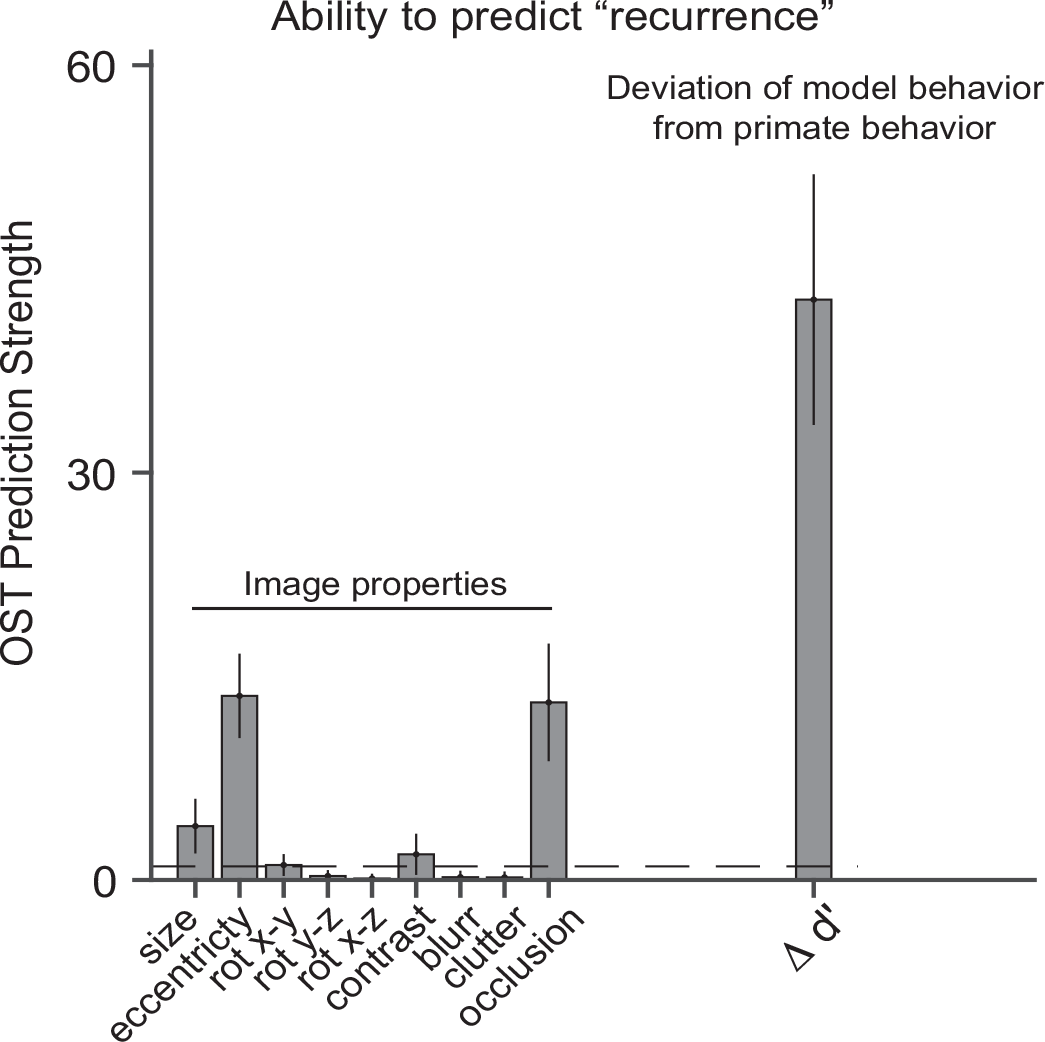
Comparison of OST prediction strength between different image properties and the ∆*d′* vector (deviation of model behavior from pooled monkey behavior). The black dashed line denotes the significance threshold of the F-statistic. Image properties like object size, eccentricity, and presence of an occluder significantly predict OST. However, the ∆*d′* vector provides the strongest OST predictions. Error bars denote the bootstrap standard deviation over images.

## Discussion

The overall goal of this study was to ask if recurrent circuits are critical to ventral stream’s execution of core recognition behavior — the ability to report object category in the central 10° with less than 200 ms of image viewing duration. We reasoned that, if computations mediated by recurrent circuits are critical for some images, then one way to find such images is by finding images that are difficult for non-recurrent DC-NNs to solve, but are nevertheless easily solved by primates. Thus we first used extensive behavioral testing to find such “challenge” images along with behaviorally matched “control” images. With these in hand, we then aimed to look for a likely empirical signature of recurrence — the requirement of additional time to complete successful processing. To ask this question, we first had to confirm that the challenge images that are behaviorally solved (by definition) were, in fact, solved by the ventral stream — as predicted by current models of the neural mechanisms underlying core recognition (Majaj et al., 2015). Using large-scale IT population neurophysiology, we confirmed part of this prediction: behaviorally-sufficient linearly decodable object solutions emerged in the IT population activity for essentially all of the challenge images (assessed with the same number of neurons and training exampled as for the control images). But looking at the temporal evolution of these IT population solutions simultaneously revealed a key observation not revealed in prior work (e.g. Majaj et al., 2015)— the IT solutions were lagged by an average of ~30ms later for challenge images compared to the “control” images. In addition, we also found that the temporally lagged IT population response patterns that contained the linearly-decodable object solutions were poorly predicted by DCNN model “neural” population responses to the same challenge images. This stands in contrast to the early IT population responses, which were much better predicted by the DCNN model, consistent with prior work (Yamins et al., 2014). Notably, we observed both of these findings during active task performance (when the animals had to report the identity of the dominant object in the image), but we found all of these results to be almost identical during passive viewing. Taken together, these results imply that automatically-evoked recurrent circuits are critical for object identification behavior even at the fast timescales of core object recognition.

The idea that “feedback” is important to vision and to object recognition is not new (see Lehky and Tanaka, 2016 for review). While broad concepts about the potential role of feedback in vision have been previously suggested and partly explored, we believe that this is the first work to examine these questions at the fast times scale of core object recognition, and the first to do so using image computable models of the neural processing to guide the choice of experiments (i.e. the images and discrimination tasks).

### Late object identity solution times in IT imply recurrent computations underlie core recognition

The most parsimonious interpretation of the results reported here is that the late phases of the stimulus evoked responses in IT depend on some type (or types) of recurrent computations that are not present in today’s non-recurrent DCNN ventral stream models. And our comparisons with behavior suggest that these IT dynamics are not epiphenomena, but are critical to the core recognition behavior. But what kind(s) of additional computations are taking place and where in the brain do those recurrent circuit elements live? We do not yet know the answers to these questions, but we can speculate to generate a testable space of hypotheses. Based on the number of synapses between V1 and IT, Tovee (1994) proposed that the ventral stream comprises of stages that are approximately 10-15 ms away from each other. Our observation of an additional processing time of 30 ms for challenge images is therefore equivalent to at least two additional processing stages. Thus, one possible hypothesis is a cortico-cortical recurrent pathway between the cortical areas including IT and lower areas like V4, V2 and V1 (similar to suggestions of Nurminen et al., 2018; Ullman, 1995; van Kerkoerle et al., 2014). This possibility is consistent with observations of temporally-specific effects in the response dynamics of V4 neurons (Fyall et al., 2017) for images with occlusion. Alternatively, the temporal lag signature we report here is also consistent with the possibility that IT is receiving important recurrent flow from downstream areas like the prefrontal and perirhinal cortices (e.g. as suggested by Bar et al., 2006). We also cannot rule out the possibility that all of the additional computations are due to recurrence within IT itself (e.g. consistent with recent models such as, Tang et al., 2017), or due to subcortical circuits (e.g. basal ganglia loops; Seger, 2008). These hypotheses are not mutually exclusive. Given all that prior work, the main contribution of our work is to take the very broad notion of “feedback” and pin down a narrower case that is both experimentally tractable (i.e. the neural phenomena is observable in IT for a prescribed set of images) and is guaranteed to have high behavioral relevance. The present results now motivate the need for direct perturbation studies that aim to independently suppress each of those circuit motifs to assess the relative importance of each of these circuit motifs. Such perturbations should be paired with IT electrophysiological recordings and behavior. The results of the present study also provide sets of images and predictions of exactly how and when IT will be disrupted when the critical circuit motif(s) is/are suppressed. Specifically, our measurements of both the ∆*d′* and the OST vectors provide observable signatures of recurrent computations that make clear predictions for such direct neural suppression studies. Based on our results here, we predict that a specific disruption of the relevant recurrent circuits will prevent the emergence of the object solutions to the challenge images in IT. This will in turn result in larger behavioral deficits in the challenge images (relative to the control images). Note however, that our results are stronger than that — they suggest exactly which set of images will be most affected (a mixture of mostly challenge images and some control images), and this knowledge can be used to optimize the image sets and behavioral tasks for these next experiments.

### Temporally specific failures of current ventral stream encoding models imply that recurrent circuits are needed to improve those models

Prior to this study, the best models of the ventral visual stream belonged to a class of feedforward DCNNs, e.g. HMO (Yamins et al., 2014), AlexNet (Krizhevsky et al., 2012) and VGG (Chatfield et al., 2014; Simonyan and Zisserman, 2014). These studies (Cadieu et al., 2014; Yamins et al., 2014) have demonstrated that feedforward DCNNs can explain ~50% of the within-animal explainable response variance in stimulus evoked V4 and IT responses (averaged responses from 70 - 170 ms post-stimulus onset). Our results here confirm that feedforward DCNNs indeed approximate ~50% of the first 30 ms (~90-120 ms) of the stimulus evoked, within-animal explainable IT response variance, thus establishing DCNNs as a good functional approximation of the feedforward pass of the primate ventral stream. However, in addition, we observed that the ability of DCNN neural populations to predict IT neural responses drops significantly at later phases of the stimulus evoked IT responses (> 150 ms after image onset, see Figure 4A). This is consistent with our inference that the late object solution times for challenge images are primarily caused by the additional processing time required by recurrent processes in the ventral stream. Recruitment of recurrent circuits in the form of both intra and inter-cortical feedback during these times might explain why the feedforward-only DCNN activations poorly predict the late IT responses.

### Unique object solution times per image motivate the search for better models of the link between IT neural population patterns and core object recognition behavior

Majaj et al. 2015 experimentally rejected a large number of alternative models that link ventral stream population activity to core object recognition behavior (aka “decoding models”). But that study was unable to reject a small set of specific models that were each quantitatively sufficient to predict the behavioral performance level for each and every tested object recognition task. That set of models was referred to as learned weighted sums of randomly selected average neuronal responses spatially distributed over monkey IT (LaWS of RAD IT). However, in the Majaj et al. (2015) study, the key predictor variable (behavioral performance) was computed as the average over all test images for that task. The authors speculated that a much finer-grain predictor variable, e.g. image-level behavioral performance, could provide a stronger test and thus might be able to falsify some or all of the LaWS of RAD IT decoding models. Here we observe that, even for images that have statistically non-distinguishable levels of behavioral performance, the linearly-decodable information in the IT population pattern varies quite substantially over the IT response time window used by many of the LaWS of RAD IT decoding models (70-170 ms post stimulus onset). Taken together, this argues that future work in this direction might successfully reject most or even all of the LaWS of RAD IT decoding models, and thus drive the field to create better neuronal-to-behavioral linking hypotheses.

### Role of recurrent computations: deliverables from these data and insights from deeper CNNs

Prior studies have strongly associated the role of recurrent computations with overcoming certain specific challenging image properties like occlusion (Spoerer et al., 2018), high levels of clutter (Walther et al., 2005) or engagement in tasks like visual pattern completion (Tang et al., 2017). While we agree that such images or task conditions might recruit recurrent processes in the ventral stream, the present work argues that this is not the most efficient approach to constrain future model development. Specifically, we have here found that a very good way to expose which images lean most of recurrent computations in the ventral stream is to find images in which the difference between feedforward-only DCNN and primate behavior (∆*d′*) is the largest. This difference is a far better predictor of the neural phenomena of recurrence than any of the text image properties (see Figure. 6).

While this is a very good way to focus experimental efforts, it does not yet expose the computational role of recurrence, i.e., the exact nature of the computational problem solved by recurrent circuits during core object recognition. Interestingly, we found that deeper CNNs like inception-v3, v4 (Szegedy et al., 2017), ResNet-50,101 (He et al., 2016), that introduce more nonlinear transformations to the image pixels, compared to shallower networks like AlexNet or VGG, are better models of the late phase of IT responses (the phase that is most behaviorally relevant for DCNN-challenge images). This is also consistent with a previous study (Liao and Poggio, 2016) where it was shown that a shallow recurrent neural network (RNN) is equivalent to a very deep CNN (e.g. ResNet) with weight sharing among the layers. Therefore, we speculate that what the computer vision community has achieved by stacking more layers into the CNNs, is a partial approximation of something that is more efficiently built into the primate brain architecture in the form of recurrent circuits. That is, during core (~200 ms) object recognition, recurrent computations act as additional non-linear transformations of the initial feedforward IT response, to produce more explicit (linearly separable) solutions. This provides a qualitative explanation for the requirement of recurrent circuits during a variety of challenging image conditions, the purpose of which is to achieve a more explicit object representation at the level of IT. What is now needed are new recurrent artificial neural networks (ANNs) that successfully incorporate these ideas. While the data presented here cannot fully specify the form of those ANNs, they will provide a strong check on any model that aims to succeed in these more advanced vision challenges where primates still outperform machines.

## ACKNOWLEDGMENTS

This research was supported by the Office of Naval Research MURI-114407 (J.J.D), and in part by the US National Eye Institute grants R01-EY014970 (J.J.D.), K99-EY022671 (E.B.I.), and the European Union’s Horizon 2020 research and innovation programme under grant agreement No 705498 (J.K.). We thank Arash Afraz for his surgical assistance.

## METHODS

### Subjects

The nonhuman subjects in our experiments were two adult male rhesus monkeys (Macaca mulatta). All human studies were done in accordance with the Massachusetts Institute of Technology Committee on the Use of Humans as Experimental Subjects. A total of 88 observers participated in the binary match to sample object discrimination task. Observers completed these 20-25 min tasks through Amazon’s Mechanical Turk, an online platform in which subjects can complete experiments for a small payment.

### Visual stimuli: generation

#### Generation of synthetic (“naturalistic”) images

High-quality images of single objects were generated using free ray-tracing software (http://www.povray.org), similar to Majaj et al. (2015). Each image consisted of a 2D projection of a 3D model (purchased from Dosch Design and TurboSquid) added to a random background. The ten objects chosen were bear, elephant, face, apple, car, dog, chair, plane, bird and zebra (Figure 1B). By varying six viewing parameters, we explored three types of identity while preserving object variation, position (x and y), rotation (x, y, and z), and size. All images were achromatic with a native resolution of 256 × 256 pixels (see Figure 1D, and Figure S1A for example images).

### Generation of natural images (photographs)

Images pertaining to the 10 nouns, were download from http://cocodataset.org. Each image was resized to 256 x 256 x3 pixel size and presented within the central 8 deg. We used the same images while testing the feedforward DCNNs.

### Quantification of image properties

We have compared the ability of different image properties to predict the putative recurrence signal, inferred from our results. These image properties were either pre-defined during the image generation process (e.g. object size, object eccentricity, and the object rotation vectors, presence of an object occluder) or computed after the image generation procedure. The post image generation properties are listed below:

Image contrast: This was defined as the variance of the luminance distribution per image (grayscale images only).
Image blur: The image processing literature contains multiple measures of image focus based on first order differentiation or smoothing followed by differentiation. We have used a technique from Santos et al. (1997) to define the focus of an image.
Image clutter: This measure (Feature Congestion) of visual clutter is related to the local variability in certain key features, e.g., color, contrast, and orientation (Rosenholtz et al., 2007).

### Primate behavioral testing

#### Humans tested on amazon mechanical turk

We measured human behavior (from 88 subjects) using the online Amazon MTurk platform which enables efficient collection of large-scale psychophysical data from crowd-sourced “human intelligence tasks” (HITs). The reliability of the online MTurk platform has been validated by comparing results obtained from online and in-lab psychophysical experiments (Majaj et al., 2015; Rajalingham et al., 2015). Each trial started with a 100 ms presentation of the sample image (one our of 1320 images). This was followed by a blank gray screen for 100 ms; followed by a choice screen with the target and distractor objects (similar to Rajalingham et al., 2018). The subjects indicated their choice by touching the screen or clicking the mouse over the target object. Each subjects saw an image only once. We collected the data such that, there were 80 unique subject responses per image, with varied distractor objects.

#### Monkeys tested during simultaneous electrophysiology

##### Active bi-nary object discrimination task

We measured monkey behavior from two male rhesus macaques. Images were presented on a 24-inch LCD monitor (1920 × 1080 at 60 Hz) positioned 42.5 cm in front of the animal. Monkeys were head fixed. Monkeys fixated a white square dot (0.2°) for 300 ms to initiate a trial. The trial started with the presentation of a sample image (from a set of 1320 images) for 100 ms. This was followed by a blank gray screen for 100 ms, after which the choice screen containing a target and a distractor object was shown. The monkey was allowed to view freely the choice images for up to 1500 ms and indicated its final choice by holding fixation over the selected image for 400 ms. Trials were aborted if gaze was not held within ±2° of the central fixation dot during any point until the choice screen was shown.

##### Passive Viewing

During the passive viewing task, monkeys fixated a white square dot (0.2°) for 300 ms to initiate a trial. We then presented a sequence of 5 to 10 images, each ON for 100 ms followed by a 100 ms gray blank screen. This was followed by a water reward and an inter trial interval of 500 ms, followed by the next sequence. Trials were aborted if gaze was not held within ±2° of the central fixation dot during any point.

#### Behavioral Metrics

We have used the same one-vs-all image level behavioral performance metric (*I*_1_) to quantify the performance of the humans, monkeys, deep HCNNs and neural based decoding models for the binary match sample tasks. This metric estimates the overall discriminability of each image containing a specific target object from all other objects (pooling across all 9 possible distractor choices). For example, given an image of object ‘*i*’, and all distractor objects (*j* ≠ *i*) we first compute the average hit rate,

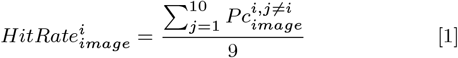

where *Pc* refers to the fraction of correct responses for the binary task between objects ‘*i*’ and ‘*j*’. We then compute the false alarm rate for the object ‘*i*’ as,

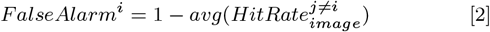

The unbiased behavioral performance, per image, was then computed using a sensitivity index *d′*,

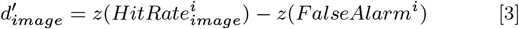
where *z* is the inverse of the cumulative Gaussian distribution. The values of *d′* were bounded between −5 and 5. Given the size of our image-set, the vector contains 1320 independent values.

### Large scale multielectrode recordings and simultaneous behavioral recording

#### Surgical implant of chronic micro-electrode arrays

Before training, we surgically implanted each monkey with a head post under aseptic conditions. After behavioral training, we recorded neural activity using 10 × 10 micro-electrode arrays (Utah arrays; Blackrock Microsystems). A total of 96 electrodes were connected per array. Each electrode was 1.5 mm long and the distance between adjacent electrodes was 400 *u*m. Before recording, we implanted each monkey multiple Utah arrays in the IT cortex (monkey M: 3 arrays in right hemisphere and two in the left hemisphere; monkey N: 3 arrays in the left hemisphere and 2 arrays in the right hemisphere). Array placement was guided by the sulcus pattern, which was visible during surgery. The electrodes were accessed through a percutaneous connector that allowed simultaneous recording from all 96 electrodes from each array. All behavioral training and testing was performed using standard operant conditioning (water reward), head stabilization, and real-time video eye tracking. All surgical and animal procedures were performed in accordance with National Institutes of Health guidelines and the Massachusetts Institute of Technology Committee on Animal Care.

#### Eye Tracking

We monitored eye movements using video eye tracking (SR Research EyeLink 1000). Using operant conditioning and water reward, our 2 subjects were trained to fixate a central white square (0.2°) within a square fixation window that ranged from ±2°. At the start of each behavioral session, monkeys performed an eye-tracking calibration task by making a saccade to a range of spatial targets and maintaining fixation for 500 ms. Calibration was repeated if drift was noticed over the course of the session.

#### Electrophysiological Recording

During each recording session, band-pass filtered (0.1 Hz to 10 kHz) neural activity was recorded continuously at a sampling rate of 20 kHz using Intan Recording Controller (Intan Technologies, LLC). The majority of the data presented here were based on multiunit activity. We detected the multiunit spikes after the raw data was collected. A multiunit spike event was defined as the threshold crossing when voltage (falling edge) deviated by less than three times the standard deviation of the raw voltage values. Of 960 implanted electrodes, five arrays (combined across the two hemispheres) × 96 electrodes × two monkeys, we focused on the 424 most visually driven, selective and reliable neural sites. Our array placements allowed us to sample neural sites from different parts of IT, along the posterior to anterior axis. However, for all the analyses, we did not consider the specific spatial location of the site, and treated each site as a random sample from a pooled IT population.

#### Neural recording quality metrics per site

Visual drive per neuron 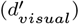: We estimated the overall visual drive for each electrode. This metric was estimated by comparing the COCO image responses of each site to a blank (gray screen) response.

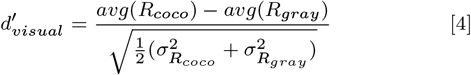

Image rank-order response reliability per neural site 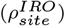: To estimate the reliability of the responses per site, we computed a spearman-brown corrected, split half (trial-based) correlation between the rank order of the image responses (all images).

Selectivity per neural site: For each site, we measured selectivity as the d’ for separating that site’s best (highest response-driving) stimulus from its worst (lowest response-driving) stimulus. *d′* was computed by comparing the response mean of the site over all trials on the best stimulus as compared to the response mean of the site over all trials on the worst stimulus, and normalized by the square-root of the mean of the variances of the sites on the two stimuli:

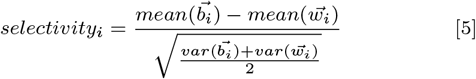

where 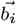 is the vector of responses of site i to its best stimulus over all trials and 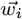 is the vector of responses of site i to its worst stimulus. We computed this number in a cross-validated fashion, picking the best and worst stimulus on a subset of trials and then computing the selectivity measure on a separate set of trials, and averaging the selectivity value of 50 trial splits.

Inclusion criterion for neural sites: For our analyses, we only included the neural recording sites that had an overall significant visual drive 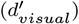, an image rank order response reliability 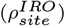 that was greater than 0.6 and a selectivity score that was greater than 1. Given that most of our neural metrics are corrected by the estimated noise at each neural site, the criterion for selection of neural sites is not that critical, and it was mostly done to reduce computation time by eliminating noisy recordings.

### Population Neural response latency estimation

Onset latencies (*t*_*onset*_) were determined as the earliest time from sample image onset when the firing rates of neurons were higher than one-tenth of the peak of its response. We averaged the latencies estimated across individual neural sites to compute the population latency. Peak latencies (*t*_*peak*_) were estimated as the time of maximum response (firing rate) of a neural site in response to an image. We averaged the peak latencies estimated across individual neural sites to compute the population peak latency per image. Both of these latency measures were computed across different sets of images (control and challenge) as mentioned in the article.

### Estimation of solution for object identity per image

#### IT cortex

To estimate what information downstream neurons could easily “read” from a given IT neural population, we used a simple, biologically plausible linear decoder (i.e., linear classifiers), that has been previously shown to link IT population activity and primate behavior (Majaj et al., 2015). Such decoders are simple in that they can perform binary classifications by computing weighted sums (each weight is analogous to the strength of synapse) of input features and separate the outputs based on a decision boundary (analogous to a neuron’s spiking threshold). Here we have used a support vector machine (SVM) algorithm with linear kernels. The SVM learning model generates a decoder with a decision boundary that is optimized to best separate images of the target object from images of the distractor objects. The optimization is done under a regularization constraint that limits the complexity of the boundary. We used L2 regularization (strength of regularization, *λ* was optimized for each train-set) and a stochastic gradient descent solver to estimate 10 (one for each object) one-vs-all classifiers. After training each of these classifiers with a set of 100 training images per object, we generated a class score (*sc*) per classifier for all held out test images. We then converted the class scores into probabilities by passing them through a softmax (normalized exponential) function.

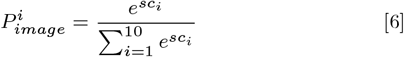

We then computed the binary task performances, by calculating the percent correct score for each pair of possible binary task given an image. For instance, if an image was from object *i*, then the percent correct score for the binary task between object *i* and object *j*, was computed as,

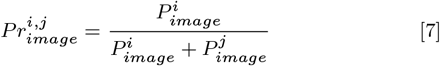

From each percent correct score, we then estimated a neural *I*_1_ score (per image), following the same procedures as the behavioral metric.

#### Object solution time per image in IT (OST_image_)

Object solution time per image, *OST_image_* was defined as the time it takes for linear IT population decodes to reach within the error margins of the pooled monkey behavioral *I*_1_ score for that image. In order to estimate this time, we first computed a neural *I*_1_ vector for non-overlapping 10 ms time bins post the sample image onset. We then used linear interpolation to predict the value of the *I*_1_ vector per image at any given time between 0 and 250 ms. We then used the Levenberg-Marquardt algorithm to estimated the time at which the neural *I*_1_ vector reached the error margins of the pooled monkey behavioral *I*_1_.

### Deep Convolutional Neural Networks (DCNN)

#### Binary object discrimination tasks with DCNNs

We used the same linear decoding scheme mentioned above for estimating the object solution strengths per image for the DCNNs. This essentially involves replacing the IT neural population features (as mentioned above) with the respective image-evoked DCNN model (e.g. AlexNet ‘fc7’ layer) features. The rest of the procedure remained the same. The features extracted from each of the models were then projected onto the first 1000 principle components (ranked in the order of variance explained) to construct the final feature set used. This was done to overcome large feature set sizes.

#### Prediction of neural response from DCNN features

We modeled each IT neural site as a linear combination of the DCNN model features. We first extracted the features per image, from the DC-NNs’ penultimate layers. The features extracted were then projected onto its first 1000 principle components (ranked in the order of variance explained) to construct the final feature set used. For example, we used the features from AlexNet’s (Krizhevsky et al., 2012) ‘fc7’ layer to generate Fig 4A. Using a 50%/50% train/test split of the images, we then estimated the regression weights (i.e how we can linearly combine the model features to predict the neural site’s responses) using a partial least squares (MATLAB command: plsregress) regression procedure, using 20 retained components. For each set of regression weights estimated on a train imageset, we generated the output of that ‘synthetic neuron’ for the held out test set. The percentage of explained variance, IT predictivity (for details refer Yamins et al., 2014) for that neural site, was then computed by normalizing the *r*^2^ prediction value for that site by the self-consistency of the image responses for that site and the self-consistency of the regression weights for that site (estimated by a Spearman Brown corrected trial split correlation score).

#### Estimation of the OST prediction strength

We compared how well different factors and ∆*d′* between monkey behavior and AlexNet ‘fc7’, predicted the differences in the object solution time (OST) estimates. Each image has an associated value for different image properties, either categorical e.g. occcluded/non-occluded or continuous e.g. object size etc. We first divided the image sets into two groups, high and low, for each factor. The high group for each factor contained images with values higher than 95th percentile of the factor distribution, and the low group contained the ones with values less than 5th percentile of the distribution. For the categorical factor like occlusion, the high group contained images with occlusion and the low group contained images without occlusion. Then, for each factor we performed a one-way ANOVA with object solution time as the dependent variable. The rationale behind this test was if the experimenter(s) were to create image sets based on any one of these factor, how likely is it expose a large difference between the OST values. Therefore, we used the F-value of the test (y-axis in Fig 6A) to quantify the OST prediction strength.

## SUPPLEMENTARY MATERIALS

The following figures have been referenced in the main article.

**Fig. S1.**
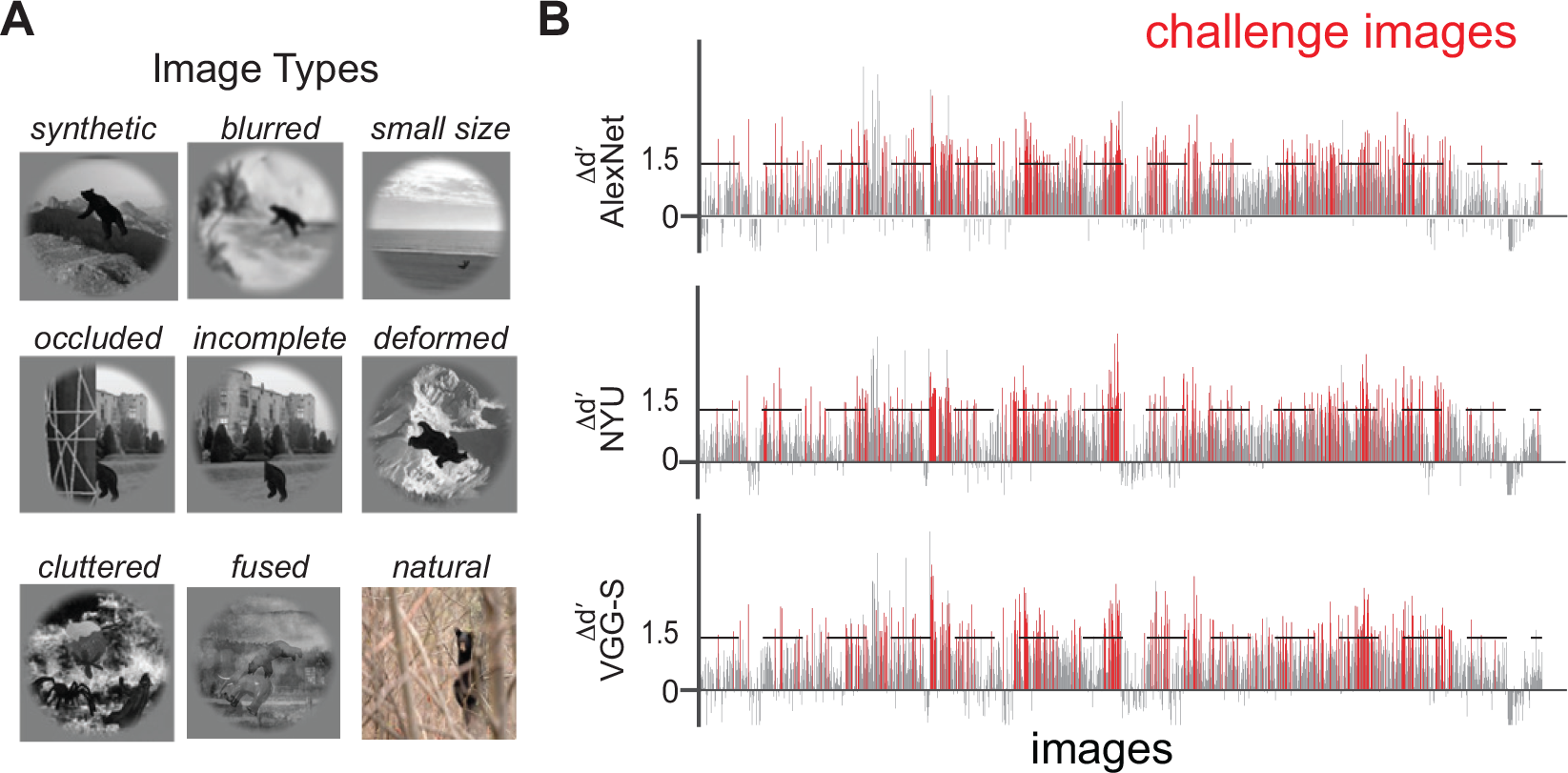
A) Examples of different image types used in the behavioral testing. Different image types included synthetic images containing an object in an uncorrelated background, images with blurr, small object sizes, occlussion, incomplete objects, deformed objects, cluttered scenes, fused objects, and natural photographs. B) Comparison of pooled monkey behavioral performance and three DCNN models with similar architecture, VGG-S (Chatfield et al., 2014; Sermanet et al., 2013), NYU (Zeiler and Fergus 2014), and AlexNet (Krizhevsky et al., 2012).

**Fig. S2.**
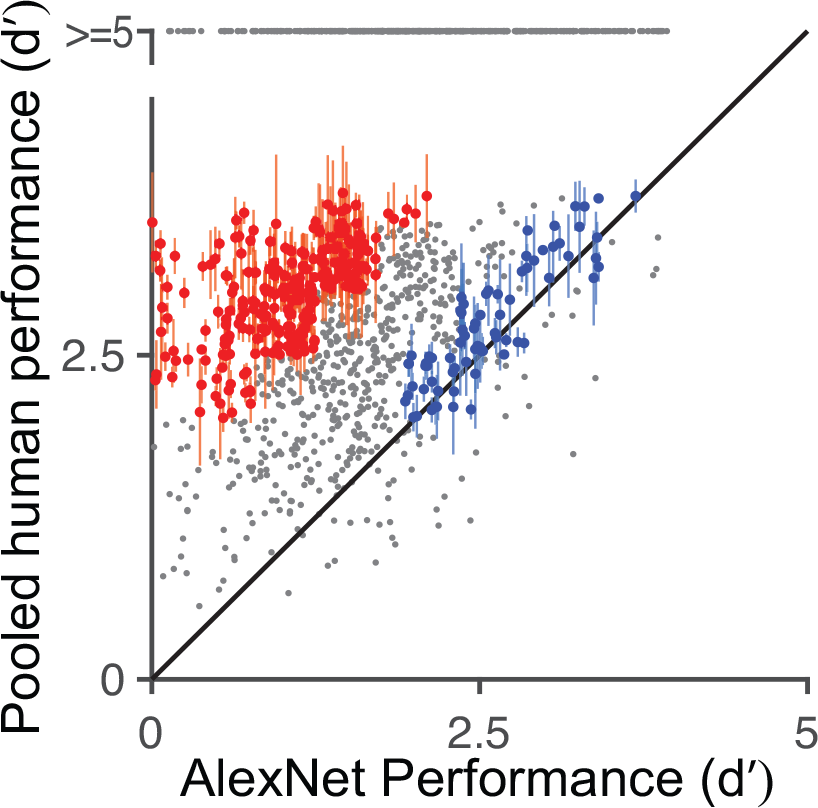
Comparison of human performance (data pooled across 88 human subjects) and DCNN performance (AlexNet; ‘fc7’ Krizhevsky et al. 2012). Each dot represents the behavioral task performance (*I*_1_; refer Methods) for a single image. We reliably identified challenge (red dots) and control (blue dots) images. Error bars are bootstrapped s.e.m.

**Fig. S3.**
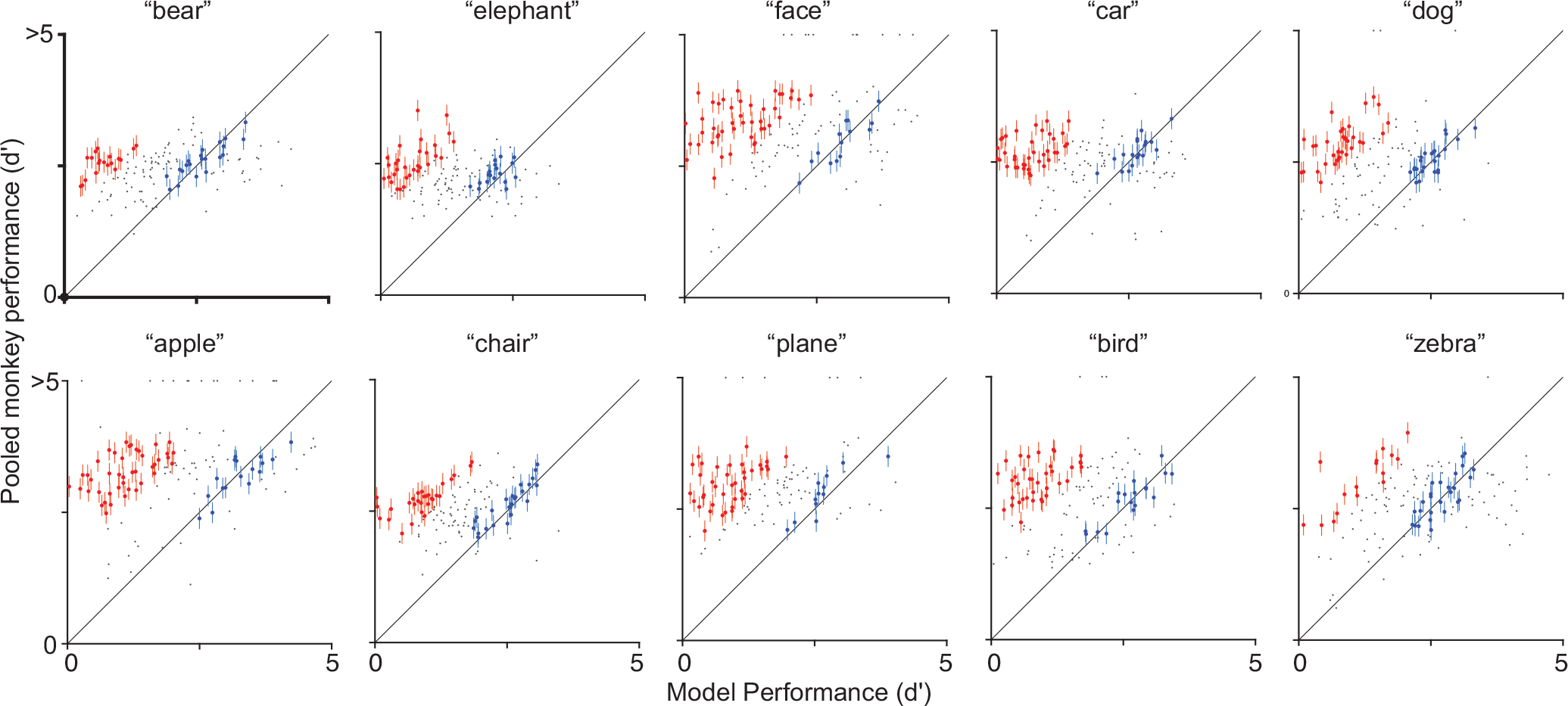
Object by object comparison of pooled monkey performance (data pooled across 2 monkeys) and DCNN performance (AlexNet; ‘fc7’ Krizhevsky et al. 2012). Each dot represents the behavioral task performance (*I*_1_; refer Methods) for a single image of the corresponding object. We reliably identified challenge (red dots) and control (blue dots) images. Error bars are bootstrapped s.e.m.

**Fig. S4.**
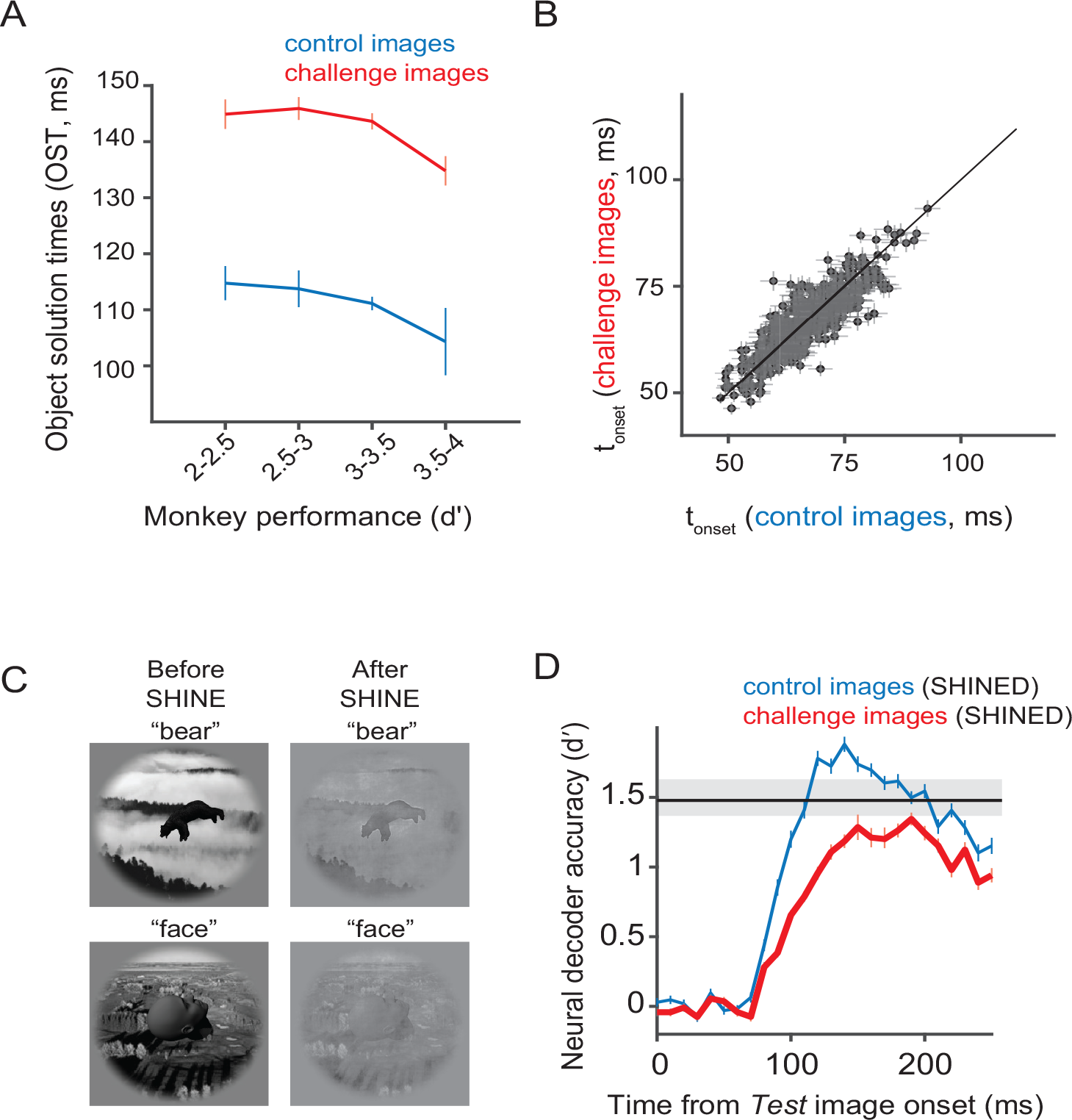
A) Dependence of *OST* on the pooled monkey *I*_1_ level. The red and the blue curves show the OST values averaged across images with behavioral *I*_1_ accuracy within the limits shown on the x-axis, for challenge and control images respectively. B) Comparison of the onset latencies (*t_onset_*) per neuron, between the challenge (y-axis) and control (x-axis) images avergaed across images of each group. Horizontal and vertical errorbars denotes s.e.m across images. C) Examples of two images, before and after the SHINE (Spectrum, histogram, and intensity normalization and equalization) algorithm was implemented. D) nAverage IT population decodes over time after the SHINE technique was implemented, for the control (blue) and challenge (red) images. The errorbars denote s.e.m across images. The black line indicates the average behavioral *I*_1_ for the pooled monkey population across all images. The gray shaded region indicates the standard deviation of the behavioral *I*_1_ for the pooled monkey population across all images

**Fig. S5.**
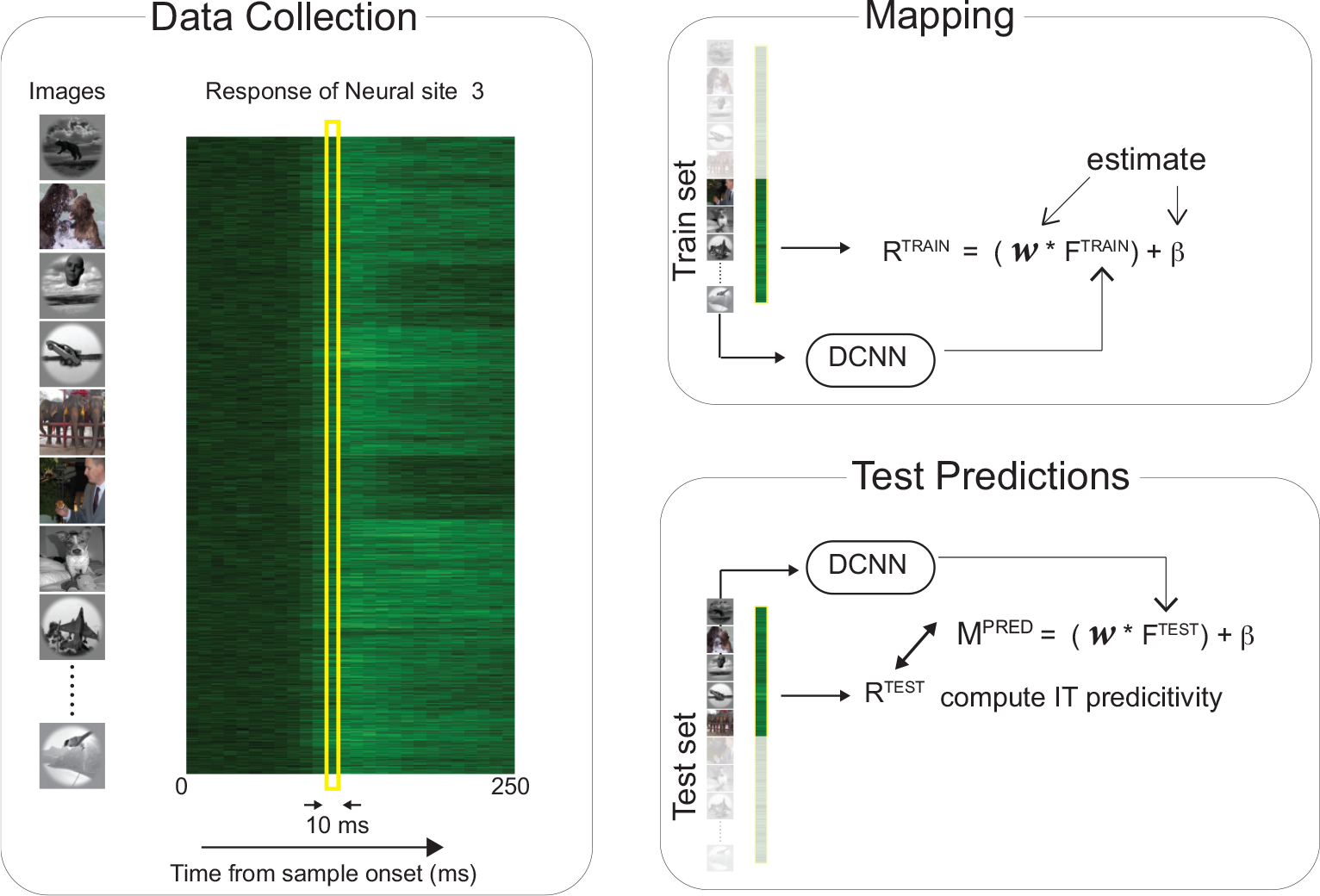
Predicting IT neural responses with DCNN features. Schematic of the DCNN neural fitting and prediction testing procedure. This includes three main steps. *Data collection*: neural responses are collected for each of the 1320 images (~50 repetitions), e.g. shown is that of neural site 3, across 10 ms timebins. *Mapping*: We divide the images and the corresponding neural features (R^TRAIN^) into a 50-50 train-test split. For the train images, we compute the image evoked activations (F^TRAIN^) of the DCNN model from a specific layer. We then use partial least square regression to estimate the set of weights (*w*) and biases (*β*) that allows us to best predict R^TRAIN^ from F^TRAIN^. *Test Predictions*: Once we have the best set of weights (*w*) and biases (*β*) that linearly map the model features onto the neural responses, we generate the predictions (M^PRED^) from this synthetic neuron for the test image evoked activations of the model FTEST. We then compare these predictions with the test image evoked neural features (R^TEST^) to compute the IT predictivity of the model.

**Fig. S6.**
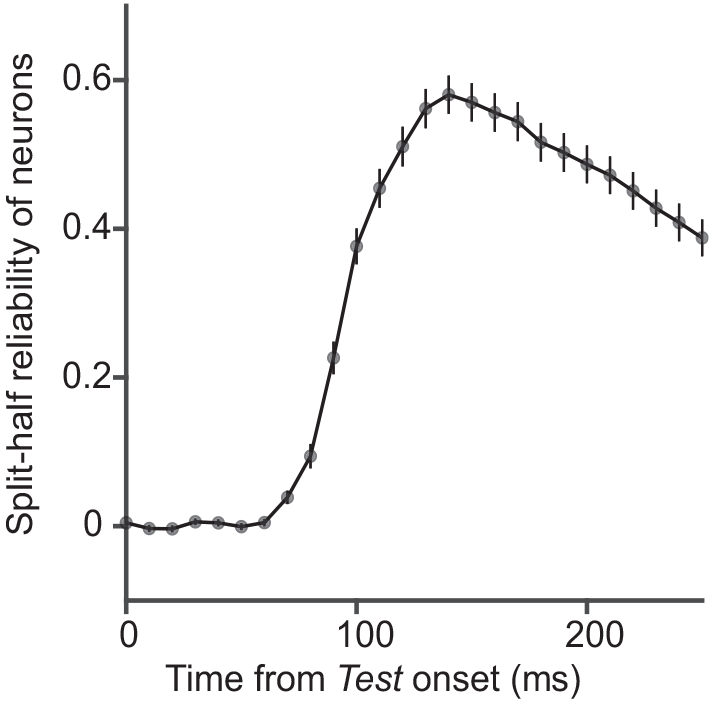
Internal consistency of the IT neural responses across time. The internal consistency was computed as a Spearman-Brown corrected correlation between two split halves (trial based) of each IT neural site’s responses across all tested images. Errorbar shows s.e.m across neural sites.

## Notes

The authors declare no competing financial interests.

